# Activity in serotonergic axons in visuomotor areas of cortex is modulated by the recent history of visuomotor coupling

**DOI:** 10.1101/2025.03.11.642559

**Authors:** Baba Yogesh, Georg B Keller

## Abstract

Visuomotor experience is necessary for the development of normal function of visual cortex (Attinger et al., 2017) and likely establishes a balance between movement-related predictions and sensory signals (Jordan and Keller, 2020). This process depends at least in part on plasticity in visual cortex (Widmer et al., 2022). Key signals involved in driving this plasticity are visuomotor prediction errors (Keller et al., 2012; Keller and Mrsic-Flogel, 2018). Ideally however, the amount of plasticity induced by an error signal should be a function of several variables - including the total prediction error across all of cortex at that moment, the animal’s experience in the current environment or task, stability of the current environment, and task engagement - for optimal computational performance. Candidates for regulators of visuomotor prediction error driven plasticity are the three major neuromodulatory systems that innervate visual cortex in the mouse: acetylcholine, noradrenaline, and serotonin. While visuomotor mismatch acutely triggers activity in noradrenaline (Jordan and Keller, 2023) but not acetylcholine (Yogesh and Keller, 2023) axons in visual cortex, how serotonergic axons in cortex respond to visuomotor mismatch is unknown. Here, we characterized the activity of serotonergic axons in visual cortex (V1) and in area A24b, a motor cortical area in anterior cingulate cortex (ACC), of awake head-fixed mice using two-photon calcium imaging. Our results reveal cortical region-specific responses to visuomotor stimuli in serotonergic axons, but no evidence of a response to visuomotor mismatch. However, average activity in serotonergic axons was modulated by the recent history of visuomotor coupling. We speculate that serotonin could function to regulate visuomotor plasticity as a function of the predictability of the environment with a slow integration time constant.

## INTRODUCTION

Visual feedback is coupled to motor output by the physical structure of bodies and the world. The brain can take advantage of this coupling to predict the sensory consequences of movement (Jordan and Rumelhart, 1992; Keller and Mrsic-Flogel, 2018). With first visuomotor experience in life, layer 2/3 neurons in visual cortex learn to combine movement related predictions and visual signals to compute visuomotor prediction errors (Attinger et al., 2017). In the fully developed circuit, movement related predictions from motor related cortical areas (Leinweber et al., 2017) are balanced against visual signals in layer 2/3 neurons (Jordan and Keller, 2020) to produce visuomotor mismatch responses. Visuomotor mismatch responses are triggered by the sudden absence of expected visual flow during locomotion in a closed loop virtual reality environment. It has been speculated that these mismatch responses could function as negative prediction errors for visuomotor learning of the internal model and also could be used to update internal representations (Keller and Mrsic-Flogel, 2018). This dual role (driving plasticity in internal models and updating internal representations) is likely the key computational advantage of a processing system based on prediction errors (see (Keller and Sterzer, 2024) for a discussion).

However, the amount of plasticity these error signals induce needs to be modulated as a function of a variety of different variables. For example, an error signal encountered during active movement needs to be interpreted differently from one experienced during passive observation. Only error signals encountered during active movement should update the internal model that produces movement-related predictions (Keller and Mrsic-Flogel, 2018). Similarly, the amount of plasticity induced by a prediction error should depend on the overall amount of concurrent prediction errors in cortex (Jordan, 2024). If a prediction error is local to only one part of visual space, the error is likely driven by an external stimulus and should only result in an update of internal representations. If, however, there is a more global error this is likely the consequence of an imperfect prediction and hence it should trigger plasticity of the internal model. Likewise, the amount of plasticity should also depend on the recent history of the level of predictability in the environment (Yu and Dayan, 2005).

Candidates for signals that could convey such plasticity modifying variables are the neuromodulatory systems that innervate visual cortex. Most prominent among these are acetylcholine, noradrenaline, and serotonin. All three neuromodulatory systems influence plasticity in cortex (Bear and Singer, 1986; Jordan and Keller, 2023; Vetencourt et al., 2011; Jitsuki et al., 2011; Froemke et al., 2007). Cholinergic axons from basal forebrain convey a locomotion-state dependent signal to V1, but exhibit no responses to visuomotor mismatch (Yogesh and Keller, 2023). Such a signal could serve as a movement state modifier of plasticity. Noradrenergic axons that project to V1 from locus coeruleus respond to visuomotor mismatch and broadcast these signals across cortex (Jordan and Keller, 2023). This could be used as a modifier of plasticity as a function of the total amount of prediction error across cortex (Jordan, 2024). Finally, the role of serotonergic signals in visuomotor integration is relatively unexplored, but given evidence from the role of serotonin in the visuomotor integration in larval zebrafish (Kawashima et al., 2016), and its role in signaling prospective value (Harkin et al., 2025), we speculated that serotonin release in V1 might act to signal the level of predictability in the environment.

The raphe nucleus in midbrain contains the majority of forebrain-projecting serotonergic neurons (Jacobs and Azmitia, 1992). In the mouse dorsal raphe, these neurons number around 9000 (Ishimura et al., 1988) and extend axonal projections throughout the entire forebrain including most of cortex (Jacobs and Azmitia, 1992; Ren et al., 2019). To investigate the potential role of serotonin in visuomotor integration in cortex, we recorded the activity of serotonergic axons in two separate regions of cortex – in A24b, a motor related area in anterior cingulate cortex that sends strong motor related input to V1, and in V1 – in awake mice using two-photon calcium imaging.

## RESULTS

To be able to record the activity of serotonergic axons in cortex using two-photon calcium imaging, we used an AAV to express an axon-targeted GCaMP6s (Broussard et al., 2018) in the dorsal raphe serotonergic neurons using SERT-Cre mice (Zhuang et al., 2005). Dorsal raphe sends diffuse axonal projections to most of the forebrain (Ren et al., 2019). We recorded calcium activity of labeled serotonergic axons in layer 1 of A24b and V1 (**Figures 1A and 1B**). Mice were head-fixed on a spherical treadmill surrounded by a toroidal screen (**Figure 1C**). We used a set of different visuomotor conditions known to activate neurons in A24b and in V1 to probe for activation of serotonergic axons. First, mice were exposed to a closed loop condition during which locomotion velocity was coupled to visual flow speed in a virtual tunnel. To probe for negative prediction error responses, we presented short (1 s) pauses in the coupling between locomotion velocity and visual flow (visuomotor mismatches) in the closed loop condition at random times. We then measured activity in an open loop condition during which the visual flow was a replay of the flow generated by the mouse in the preceding closed loop condition and thus uncoupled from locomotion. Finally, we measured activity during the presentation of full field drifting gratings. Throughout all experimental conditions mice were free to locomote on the spherical treadmill.

**Figure 1.**
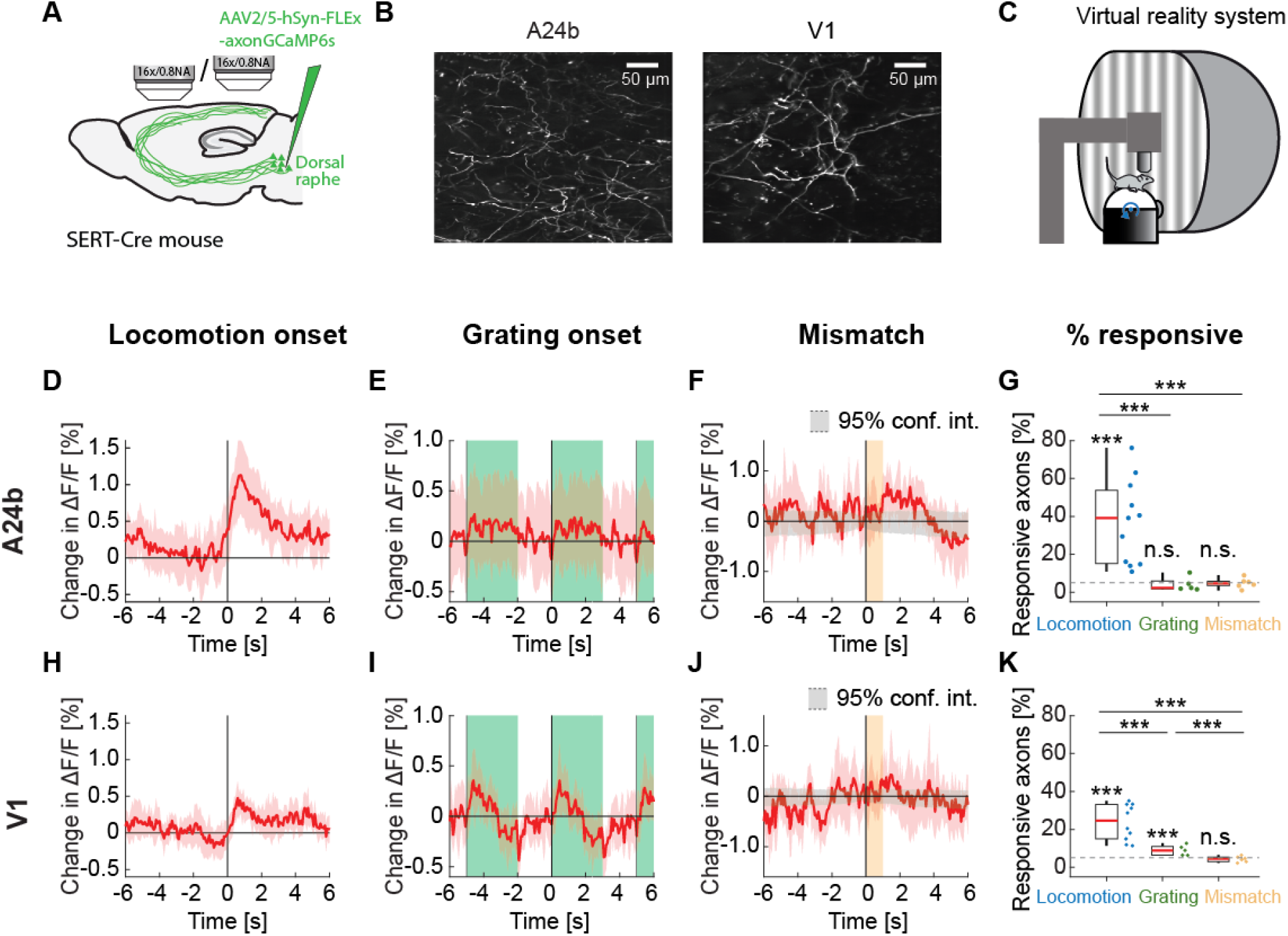
Serotonergic axon activity in A24b and V1. **(A)** We injected an AAV virus into dorsal raphe of SERT-Cre mice to express axonal GCaMP6s in serotonergic neurons. We performed two-photon calcium imaging of the labelled serotonergic axons in A24b and in V1. **(B)** Example two-photon image of serotonergic axons in A24b (left) and in V1 (right). **(C)** Schematic of the virtual reality setup. Mice were head-fixed on a spherical treadmill surrounded by a toroidal screen on which visual stimuli were presented in different visuomotor conditions. **(D)** Average response of serotonergic axons to locomotion onset in A24b. Mean (solid lines) and the bootstrap error indicating 95% confidence interval (shading) are calculated as hierarchical bootstrap estimate for each time bin. **(E)** As in **D**, but for grating onsets in A24b. Green shading marks the duration of grating presentation. **(F)** As in **D**, but for visuomotor mismatch onsets in A24b. Orange shading marks the duration of visuomotor mismatch. As mismatch events occur only during times of locomotion and locomotion itself drives activity in these axons, we quantified the distribution of serotonergic axon activity on random triggers during locomotion (95% confidence interval, gray shading). **(G)** The fraction of serotonergic axons in A24b responsive to locomotion, grating, and visuomotor mismatch onset, quantified for each imaging site. Each datapoint is one imaging site. Boxes show 25^th^ and 75^th^ percentile, central mark is the median, and the whiskers extend to the most extreme data points not considered outliers. Dashed line marks chance level. n.s.: not significant; *p<0.05; **p<0.01; ***p<0.001; see **Table S1** for statistical information. **(H)** As in **D**, but for serotonergic axons in V1. **(I)** As in **E**, but for serotonergic axons in V1. **(J)** As in **F**, but for serotonergic axons in V1. **(K)** As in **G**, but for serotonergic axons in V1.

On locomotion onset, we found that the activity in many serotonergic axons increased in both A24b and V1 (**Figure S1**). However, different from cholinergic activity that is sustained during locomotion (Yogesh and Keller, 2023), the activity increases were primarily transient (**Figure 1D**). We found no evidence of a response in serotonergic axons in A24b to the presentation of grating stimuli (**Figure 1E**) or to visuomotor mismatch (**Figure 1F**). This was reflected in the fraction of responsive serotonergic axons, with about 40% of the imaged axons in A24b being significantly responsive to locomotion onset, while the fraction of responsive axons was at chance for grating and visuomotor mismatch onsets (**Figure 1G**). Interestingly, the activity of serotonergic axons in V1 exhibited a slightly different pattern. In V1, serotonergic axons increased their activity on locomotion onset (**Figure 1H**), but less strongly than in A24b. In addition, in V1 these axons exhibited clear grating responses (**Figure 1I**). Looking at the difference between locomotion onset response and grating response illustrates the region specificity of the responses (**Figure S2**). But as in A24b, we found no evidence of a response to visuomotor mismatch in V1 (**Figure 1J**). This response profile was again reflected in the fraction of responsive axons, being significant for locomotion and grating onsets, but at chance for visuomotor mismatch (**Figure 1K**). Thus, similar to findings in zebra fish (Mutlu et al., 2025), we find that the activity of serotonergic axons appears to be different for different target regions. This could be explained by a high covariance between serotonergic axons and neuronal activity in the target region (Mutlu et al., 2025).

Activity in serotonergic neurons can drive changes in pupil size (Cazettes et al., 2021). Given that locomotion and pupil size also correlate, we investigated whether locomotion or pupil size was a better correlate of serotonergic activity. We did this by comparing the correlation of activity in serotonergic axons with locomotion velocity to that with pupil diameter. Again, different from cholinergic activity in V1, we found that the activity of these axons in both A24b and V1 was more strongly correlated with pupil size than with locomotion velocity (**Figure 2**). Thus, given that pupil dilation is a correlate of arousal, this would be consistent with the idea that activity in serotonin axons in cortex is influenced by a process related to arousal.

**Figure 2.**
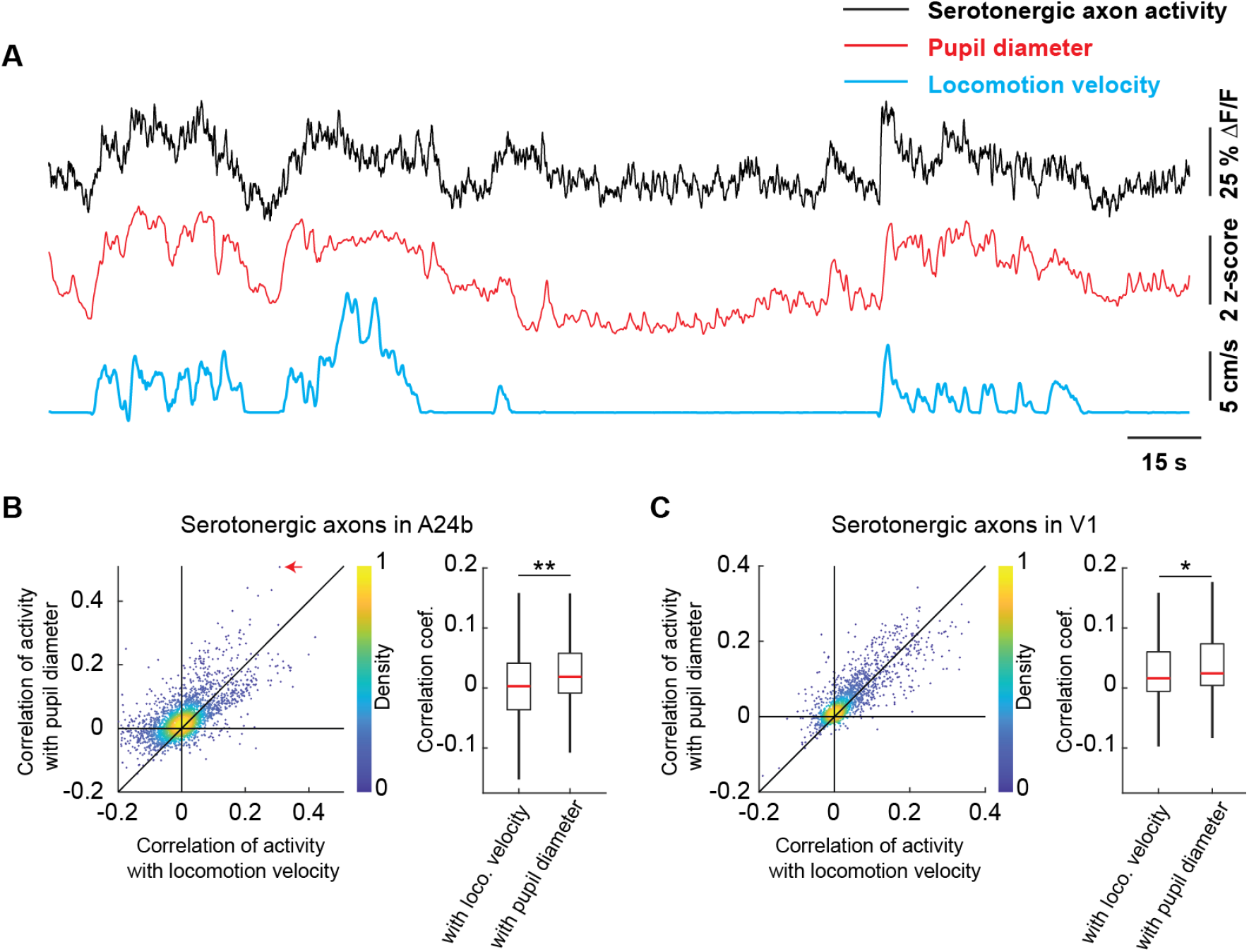
Serotonergic axon activity is better correlated with pupil diameter than with locomotion velocity. **(A)** Example calcium response trace (black) of a serotonergic axon in A24b with corresponding pupil diameter (red) and locomotion velocity (blue) traces. **(B)** Left: For all serotonergic axons in A24b, the correlation of their calcium activity with locomotion velocity plotted against the correlation with pupil diameter. Red arrow indicates the example axon shown in **A**. Right: Distributions of correlation coefficients between calcium activity and locomotion velocity, and calcium activity and pupil diameter. Same data as shown on the left. Boxes show 25th and 75th percentile, central mark is the median, and the whiskers extend to the most extreme data points not considered outliers. n.s.: not significant; *p<0.05; **p<0.01; ***p<0.001; see **Table S1** for statistical information. **(C)** As in **B**, but for serotonergic axons in V1.

To test if the recent history of visuomotor prediction errors also influences activity of serotonergic axons in cortex, we compared their activity in a visuomotor condition with high rate of visuomotor prediction errors (open loop) to a condition with lower rates of visuomotor prediction errors (closed loop). In the open loop session, visual feedback is not predictable from motor output. On first approximation, this should drive frequent visuomotor prediction errors. In the closed loop session, visuomotor prediction errors are primarily driven by isolated visuomotor mismatch events, and technical imperfections in the virtual reality system used to couple movement and visual feedback. Consistent with this idea, we found that the activity of serotonergic axons was higher in the open loop condition than in the closed loop condition in V1 (**Figure 3B**). While this was also true in A24b, the effect was not significant here (**Figure 3A**). To avoid confounding the analysis by movement or visual flow we restricted the analysis to times when the mouse was stationary and there was no visual flow.

**Figure 3.**
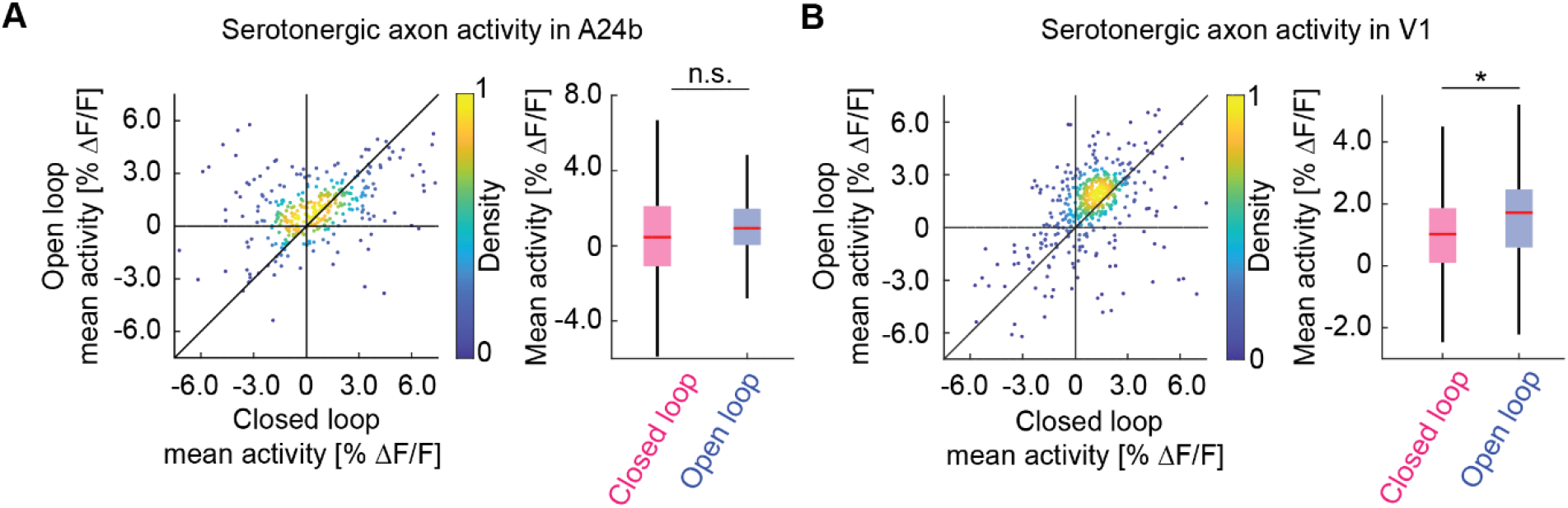
Serotonergic axon activity in V1 is higher in open loop than in closed loop conditions. **(A)** Left: For serotonergic axons in A24b, the mean calcium activity while the mice were stationary in closed loop plotted against the same while mice were stationary and there was no visual flow in open loop. For all analysis shown in this figure, we only included the 50% most active axons. Right: Boxes show 25th and 75th percentile, central mark is the median, and the whiskers extend to the most extreme data points not considered outliers. n.s.: not significant; *p<0.05; **p<0.01; ***p<0.001; see **Table S1** for statistical information. **(B)** As in **A**, but for serotonergic axons in V1.

Finally, in larval zebrafish the activity of serotonergic dorsal raphe neurons scales with the gain of visuomotor coupling (Kawashima et al., 2016). To test whether this was also the case for the activity in serotonergic axons in cortex, we adapted the stochastic gain paradigm of that work by changing the gain of visuomotor coupling in blocks of either low, medium, or high gain in random sequence (**Methods**). Consistent with the findings in larval zebrafish, we found serotonergic axon activity to be higher in high gain visuomotor blocks than in low gain visuomotor block in both A24b and V1 (**Figure 4**). Here again the analysis was restricted to times when the mouse was stationary and there was no visual flow. Thus, we conclude that the recent history of visuomotor coupling systematically modulates serotonergic activity in V1 and A24b.

**Figure 4.**
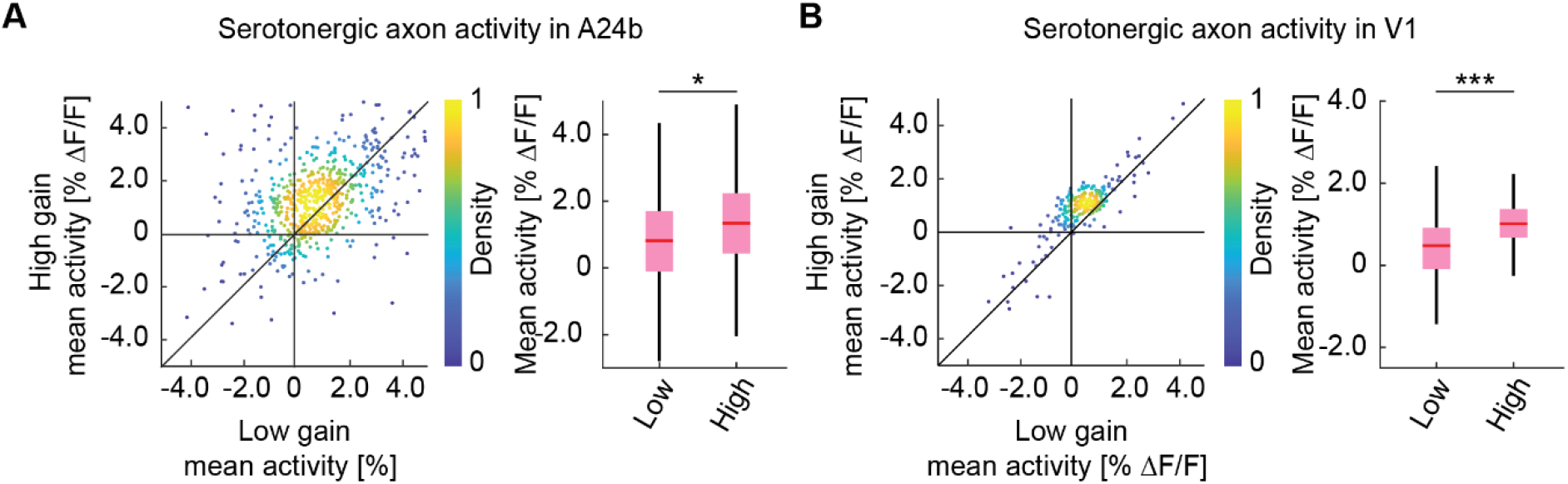
Serotonergic axon activity increases with increasing visuomotor gain. **(A)** Left: For serotonergic axons in A24b, the mean calcium activity of individual axons while the mice were in closed loop during low visuomotor gain condition plotted against the same during high visuomotor gain. As in Figure 3, we use only times when mice where stationary and there was no visual flow. Right: Boxes show 25th and 75th percentile, central mark is the median, and the whiskers extend to the most extreme data points not considered outliers. n.s.: not significant; *p<0.05; **p<0.01; ***p<0.001; see **Table S1** for statistical information. **(B)** As in **A**, but for mean calcium activity of serotonergic axons in V1.

## DISCUSSION

Traditionally, the serotonergic system has been implicated in processing of reward and punishment (Amo et al., 2014; Maswood et al., 1998; Nakamura et al., 2008), signaling the global reward state (Daw et al., 2002), and prolonging the duration of waiting behavior for delayed rewards (Fonseca et al., 2015; Miyazaki et al., 2011, 2014). An alternate line of inquiry has shown that serotonin also functions to regulate cognitive flexibility in a changing environment (Bari et al., 2010; Clarke et al., 2004; Matias et al., 2017), and can be modulated by uncertainty in the environment (Grossman et al., 2020) and signals a prospective code for value (Harkin et al., 2025). This is consistent with the idea that serotonin functions to regulate learning rates in cortex.

Whether serotonin could play a similar role of modulating visuomotor learning rates in cortex was still unclear. Previous attempts at measuring activity in serotonergic axons in visual cortex found no locomotion or pupil related activity responses (Larsen et al., 2018). In our experiments, we found clear responses to locomotion onsets and visual stimuli. Moreover, consistent with target specific activity of serotonergic axons in zebra fish (Mutlu et al., 2025), we found that axonal responses appear to be related to the coding space of the target cortical regions with stronger visual responses in serotonergic axons projecting to visual cortex and stronger locomotion onset responses in axons projecting to A24b.

One caveat of our data we cannot exclude is an influence of hemodynamic occlusion (Yogesh et al., 2025). The primary driver of hemodynamic changes are locomotion and sensory stimulation. Hence, the locomotion and grating responses we find could be contaminated by hemodynamic signals. Given that serotonergic axons we recorded were in layer 1, we expect to find less influence of occlusion effects. Finally, given that we find no evidence of hemodynamic differences between closed loop and open loop running onsets (Yogesh et al., 2025), we don’t think the differences in quiescent phases of open loop and closed loop sessions we find here could be explained by hemodynamic differences.

The finding that serotonergic axon activity was better explained by pupil diameter than by locomotion velocity indicates that the primary driver of serotonergic activity in cortex is not movement state, but something related to arousal more generally. One such driver of arousal could be uncertainty in the environment. Activity in serotonergic neurons in the dorsal raphe is modulated by an accumulation of recent reward prediction errors. This can be thought of as expected uncertainty in the environment (Grossman et al., 2020). In our paradigm, the open loop condition triggers an increase in the frequency of visuomotor prediction errors and likely results in an increase of uncertainty in the environment. Alternatively, if we assume that being in control of the environment is rewarding, the cessation of an open loop experience (to avoid confounds of locomotion and visual flow, we restrict analysis to sitting periods during which the mouse in principle is in a closed loop condition – no locomotion, no visual flow) could be interpreted as a rebound from negative value in a prospective value interpretation of serotonergic activity (Harkin et al., 2025). Consistent with either, we see higher activity in serotonergic axons in V1 during the open loop condition. In combination with the fact that we find no evidence of a response to visuomotor mismatch, these results are consistent with the interpretation that serotonergic axons in visuomotor areas of cortex signal the predictability of visuomotor coupling on long, but not short, timescales.

It has also been shown that serotonergic neurons increase their activity in conditions of increased sensorimotor gain (Kawashima et al., 2016). This could be the result of the serotonergic system tracking the gain of the visuomotor coupling, or a consequence of the fact that higher visuomotor gains in virtual reality environments are associated with a larger relative impact of (technical or biological) delays in visuomotor coupling. Thus, gain and uncertainty could be related in the sense that high gain results in lower predictability. Either way, consistent with the results in larval zebrafish (Kawashima et al., 2016), we find that the activity in serotonergic axons in A24b and V1 increase if the mouse recently experienced higher visuomotor gain.

Thus, we speculate that the three main neuromodulatory systems innervating visual cortex orchestrate to regulate different aspects of visuomotor plasticity. Cholinergic input to gate plasticity to visuomotor circuits during times of movement (Yogesh and Keller, 2023). Noradrenergic input to signal global model failure (Jordan, 2024). And serotonergic input to signal the predictability of the current environment.

## ACKNOWLEDGEMENTS

We thank all the members of the Keller lab for discussion and support. This project has received funding from the Swiss National Science Foundation (GBK), the Novartis Research Foundation (GBK), and the European Research Council (ERC) under the European Union’s Horizon 2020 research and innovation programme (grant agreement No 865617) (GBK). A preprint version of this article has been peer-reviewed and recommended by PCI Neuro (https://doi.org/10.24072/pci.neuro.100223).

## AUTHOR CONTRIBUTIONS

BY designed and performed the experiments and analyzed the data. All authors wrote the manuscript.

## METHODS

### Mice

All mice used in these experiments were SERT-Cre (Zhuang et al., 2005) heterozygotes, kept on a C57BL/6 background. A total of 4 mice, both male and female, 6-16 weeks old at the start of the experiment, were used. Between experiments, mice were group-housed in a vivarium (light/dark cycle: 12/12 hours). All animal procedures were approved by and carried out in accordance with the guidelines laid by the Veterinary Department of the Canton of Basel-Stadt, Switzerland.

### Surgery

For all surgical procedures, mice were anesthetized with a mixture of fentanyl (0.05 mg/kg; Actavis), midazolam (5.0 mg/kg; Dormicum, Roche), and medetomidine (0.5 mg/kg; Domitor, Orion) injected intraperitoneally. Analgesics were applied perioperatively (2% lidocaine gel, meloxicam 5 mg/kg) and postoperatively (buprenorphine 0.1 mg/kg, meloxicam 5 mg/kg). Eyes were covered with ophthalmic gel (Virbac Schweiz AG). Cranial windows were implanted over V1 and A24b as previously described (Keller et al., 2012; Leinweber et al., 2014). Briefly, using a dental drill, a 4 mm craniotomy was made over the right visual cortex, centered 2.5 mm lateral and 0.5 mm anterior to lambda. A second craniotomy was made over right A24b, centered at midline, 0.5 mm anterior to bregma. The exposed cortex was sealed with a 4 mm circular glass coverslip and glued in place using gel superglue (Ultra Gel, Pattex). The remaining exposed surface of the skull was covered with Histoacryl (B. Braun), and a titanium head bar was fixed to the skull using dental cement (Paladur, Heraeus Kulzer). After surgery, anesthesia was antagonized by a mixture of flumazenil (0.5 mg/kg; Anexate, Roche) and atipamezole (2.5 mg/kg; Antisedan, Orion Pharma) injected intraperitoneally.

### Axonal labeling

To image the activity of dorsal raphe serotonergic axons in A24b and V1, we expressed a calcium indicator in dorsal raphe serotonin neurons. Surgery was performed as described above, and AAV2/5-hSyn1-FLEx-axon-GCaMP6s (10^13^ GC/ml) virus was injected ipsilateral to the recording site at coordinates (AP, ML, DV relative to lambda (in mm): 0, 0, −2.3; −0.4, 0, −2.3; −0.4, 0, −2.0; −0.4, 0, −1.8; −0.8, 0, −2.3) in SERT-Cre mice to target dorsal raphe.

### Virtual reality environment

The virtual reality setup is based on the design of Dombeck and colleagues (Dombeck et al., 2007). Briefly, mice were head-fixed and free to run on an air-supported spherical treadmill. The rotation of the ball was restricted around the vertical axis with a pin. The virtual reality environment was projected onto a toroidal screen covering the mouse’s visual field approximately 240 degrees horizontally and 100 degrees vertically, using a projector (Samsung SP-F10M) synchronized to the resonant scanner of the two-photon microscope. The virtual environment consisted of an infinite corridor with walls patterned with vertical sinusoidal gratings with a spatial frequency of approximately 0.04 cycles per degree (Leinweber et al., 2014). In closed loop sessions, the visual flow speed was coupled to locomotion velocity of mice along a virtual tunnel. The gain of coupling was set such that the temporal frequency of the grating on the tunnel walls was roughly 2Hz at average running speed (across mice). In open loop sessions, we uncoupled the two and replayed the visual flow from a preceding closed loop session. In grating sessions, we presented full field drifting gratings (0°, 45°, 90°, 270°, moving in either direction) in a pseudo-random sequence. Grating stimuli were presented for 3 s. In the inter-stimulus interval (2 s), mice were shown a gray screen with average luminance matched to that of the grating stimuli.

### Stochastic gain paradigm

To test if serotonergic axon activity scales with visuomotor gain, the gain between locomotion velocity and the resultant visual flow speed was pseudo-randomly set to a factor of either 0.2 (low gain), 0.4 (medium gain) or 0.6 (high gain), in blocks of either 30 s or 60 s. These sessions were conducted on separate days from the ones used to characterize the response properties of serotonergic axons.

### Eye tracking

During all experiments, we recorded the mouse’s left eye (contralateral to the imaged hemisphere) with a CMOS infrared camera at 30 Hz frame rate. The pupil was backlit by the 930 nm laser used for two-photon imaging. We calculated pupil diameter offline by fitting a circle to the pupil. Frames with occluded pupil (during blinks and grooming) were excluded from analysis.

### Two-photon microscopy

Functional two-photon calcium imaging was performed using custom-built two-photon microscopes (Leinweber et al., 2014). The illumination source was a tunable femtosecond laser (Insight, Spectra Physics or Chameleon, Coherent) tuned to 930 nm. Emission light was band-pass filtered using a 525/50 filter for GCaMP and detected using a GaAsP photomultiplier (H7422, Hamamatsu). Photomultiplier signals were amplified (DHPCA-100, Femto), digitized (NI5772, National Instruments) at 800 MHz, and band-pass filtered at 80 MHz using a digital Fourier-transform filter implemented in custom-written software on an FPGA (NI5772, National Instruments). The scanning system of the microscopes was based on a 12 kHz resonant scanner (Cambridge Technology). Images were acquired at a resolution of 750 x 400 pixels (60 Hz frame rate), and a piezo-electric linear actuator (P-726, Physik Instrumente) was used to move the objective (Nikon 16x, 0.8 NA) in steps of 15 µm between frames to acquire images at 4 different depths. This resulted in an effective frame rate of 15 Hz. The field of view was 375 µm x 300 µm.

### Extraction of neuronal activity

Calcium imaging data were processed as previously described (Keller et al., 2012) and all data analysis was done in MATLAB (MathWorks). Briefly, raw images were full-frame registered to correct for lateral brain motion. Axons were manually selected based on mean and maximum fluorescence images. We did not merge different axonal segments based on activity similarity, as is sometimes done for cortical axon imaging (Leinweber et al., 2017). Correlation of activity between axons was too high for us to be able to identify a clear cutoff in the similarity between segments on the same axon, and on different axons. Thus we are overestimating both the total number of axons in our dataset. Raw fluorescence traces were corrected for slow drift in fluorescence using an 8^th^-percentile filtering with a 66 s (or 1000 frames) window (Dombeck et al., 2007). ΔF/F traces were calculated as mean fluorescence in a selected region of every imaging frame, subtracted and normalized by the overall median fluorescence. All neuronal calcium activity data was acquired at 15 Hz.

### Data analysis

No power analysis was performed prior to conducting the study. All data analysis was done using custom scripts written in MATLAB (MathWorks). To quantify the average population response traces, we first calculated the average event-triggered fluorescence trace for each region of interest (ROI). The responses of all ROIs were then averaged and baseline-subtracted.

Locomotion onset was defined as locomotion velocity crossing a threshold of 0.25 cm/s for at least 1 s, while having been below the threshold for 1 s before. Visuomotor mismatch responses were probed by presenting brief 1 s full field visual flow halts in the closed loop condition. For a mismatch event to be included in analysis, mice had to be locomoting uninterrupted above threshold (0.25 cm/s) from −0.5 s to +1 s after the event onset. Additionally, for a ROI to be included for analysis of the response to a particular event, we had to have data for at least 3 onsets to the event. The calcium responses were baseline subtracted using a −0.5 s to 0 s window relative to onset for response to grating and visuomotor mismatch, or −2 s to −1 s for locomotion onset. In **Figure 1G and 1K**, for each axon, we averaged the response from +0.5 s to +1.5 s, with a baseline subtraction window of −1.0 s to −0.5 s, relative to onset for individual grating onset and visuomotor mismatch trials and tested if the mean of this distribution is significantly different from 0 using a t-test with a chance threshold of 5%. The response window for locomotion onset was +0.5 s to +2.0 s. The traces in **Figure 2A** are smoothed with a sliding window of 5 frames. For most experiments we recorded multiple sequences of closed and open loop sessions. For the analysis shown in **Figure 3**, we used only data from the first closed loop and open loop presentation to avoid any confounding effects of recent experience (other than the mice normally experiencing closed loop coupling). The mice were acclimatized to the setup and got used to running on the treadmill over 5-10 hours. This was done in the dark. All recordings from a given site happened in a single session. Of the 10 such recordings that went into **Figure 3**, in 8 recordings the first two consecutive sessions, in order, were closed loop and open loop, and in the remaining 2, they were open loop and then closed loop. For the stochastic gain paradigm (**Figure 4**), the recording was performed on a different day from a different site, and mice were exposed exclusively to closed loop in that session, albeit with varying visuomotor gain. For **Figures 3 and 4**, to avoid the influence of the change in locomotion state, we excluded the first and last 2 s of each stationary bout from analysis, and included only those stationary bouts which after padding were at least 5 s long (or 3 s in case of **Figure 4** due to insufficient triggers). Recordings with less than a total of 15 s of stationary period in either of the compared conditions, were excluded from **Figures 3 and 4** (1 of 19).

### Statistical Analysis

All statistical information for the tests performed in the manuscript is provided in **Table S1**. Unless stated otherwise, the shading indicates the 95% confidence interval. For analysis where the experimental unit was axons or imaging sites (see **Table S1**), we used hierarchical bootstrap (Saravanan et al., 2020) for statistical testing due to the nested nature (axons or imaging sites and mice) of the data. Briefly, we first resampled the data (with replacement) at the level of imaging sites, and then, from the selected sites, resampled for axons. The unit (axons or sites) over which testing was done are boldfaced in the table. We then computed the mean (or median) of this bootstrap sample and repeated this N times to generate a bootstrap distribution of the mean estimate. For all statistical testing the number of bootstrap samples (N) was 10 000, for plotting bootstrap mean and standard error response curves it was 1000. The bootstrap standard error is the 95% confidence interval (2.5^th^ percentile to 97.5^th^ percentile) in the bootstrap distribution of the mean.

## Key Resources Table

| Reagent type (species) or resource | Designation | Source or reference | Identifier | Additional information |
| --- | --- | --- | --- | --- |
| Strain, strain background (adeno-associated virus) | AAV2/5-hSyn1-FLEX-axon-GCaMP6s (10<sup>13</sup> GC/ml) | FMI vector core | vector.fmi.ch | - |
| Chemical compound, drug | Fentanyl citrate | Actavis | CAS 990-73-8 | Anesthetic compound |
| Chemical compound, drug | Midazolam (Dormicum) | Roche | CAS 59467-96-8 | Anesthetic compound |
| Chemical compound, drug | Medetomidine (Domitor) | Orion Pharma | CAS 86347-14-0 | Anesthetic compound |
| Chemical compound, drug | Ropivacaine | Presenius Kabi | CAS 132112-35-7 | Analgesic compound |
| Chemical compound, drug | Lidocaine | Bichsel | CAS 137-58-6 | Analgesic compound |
| Chemical compound, drug | Buprenorphine | Reckitt Benckiser Healthcare | CAS 52485-79-7 | Analgesic compound |
| Chemical compound, drug | Humigel | Virbac | - | Ophthalmic gel |
| Chemical compound, drug | Flumazenil (Anexate) | Roche | CAS 78755-81-4 | Anesthetic antagonist |
| Chemical compound, drug | Atipamezole (Antisedan) | Orion Pharma | CAS 104054-27-5 | Anesthetic antagonist |
| Chemical compound, drug | Metacam | Boehringer Ingelheim | CAS 71125-39-8 | Analgesic compound |
| Chemical compound, drug | N-Butyl-2-cyanoacrylate | Braun | CAS 6606-65-1 | Histoacryl |
| Chemical compound, drug | Dental cement (Paladur) | Heraeus Kulzer | CAS 9066-86-8 |  |
| Genetic reagent<br>*Mus musculus* | B6.129(Cg)-Slc6a4<sup>tm1(cre)Xz</sup>/J<br>Alias used here: SERT-Cre | Jackson Laboratories | RRID:IMSR_JAX:028861 | Cre expression in serotonergic neurons |
| Software, algorithm | MATLAB (2023b) | The MathWorks | RRID:SCR_001622 | Data analysis |
| Software, algorithm | LabVIEW | National Instruments | RRID:SCR_014325 | Hardware control |
| Software, algorithm | Two-photon acquisition software | Keller laboratory | sourceforge.net/p/iris-scanning/ | Data acquisition |
| Software, algorithm | Image data processing software | Keller laboratory | sourceforge.net/p/iris-scanning/calliope | Data processing |
| Software, algorithm | Python | python.org | RRID:SCR_008394 | Virtual reality |
| Software, algorithm | Panda3D | panda3d.org | N/A | Virtual reality |
| Other | Virtual reality and two-photon setup | (Leinweber et al., 2014, 2017) | DOI: 10.3791/50885, 10.1016/j.neuron.2017.08.036 | Hardware setup |
| Other | Titanium headplate | FMI/ETHZ workshop | N/A | Mice head-fixation |
| Other | Dental drill | Meisinger | N/A | For craniotomy |

## Supplementary information

**Figure S1.**
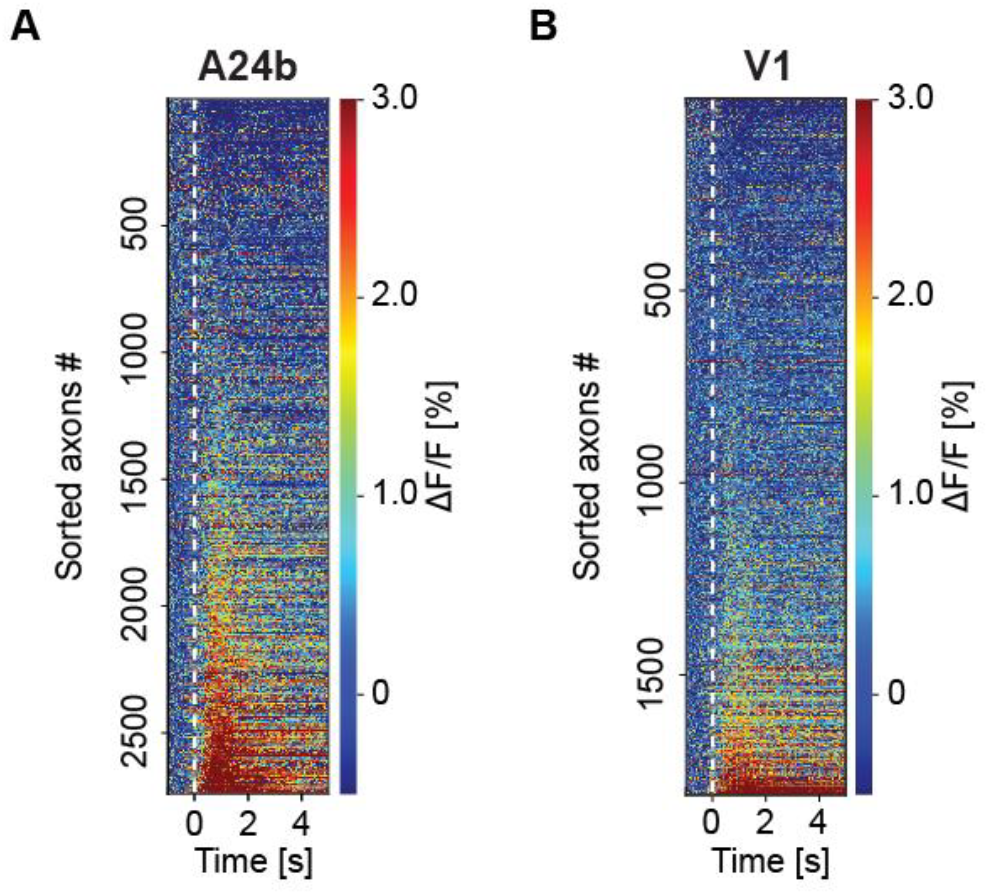
Serotonergic axons were active on locomotion onset. **(A)** Average locomotion onset activity of all serotonergic axons in A24b, sorted by their average response during locomotion onset. Please note, shown are all data and sorting introduces regression to the mean artefacts, like the apparent decrease in a subset of axons on running onset. **(B)** As in **A**, but for serotonergic axons in V1.

**Figure S2.**
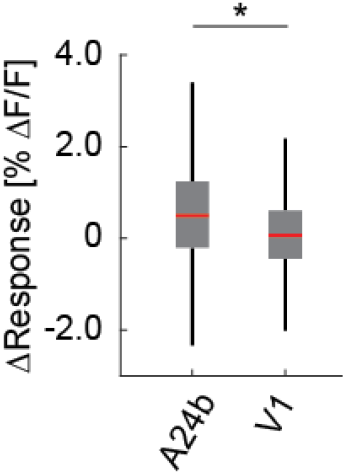
Responses of serotonergic axons were area specific. The difference in the average response on locomotion onset to grating onset for individual serotonergic axons in A24b was higher than in V1. Boxes show 25th and 75th percentile, central mark is the median, and the whiskers extend to the most extreme data points not considered outliers. n.s.: not significant; *p<0.05; **p<0.01; ***p<0.001; see **Table S1** for statistical information.

**Table S1.** Statistical information on all analysis.

| Figure panel | Value compared | Test | P-value | N (ROIs) | N (Sites) | N (Mice) |
| --- | --- | --- | --- | --- | --- | --- |
| **1D** | Locomotion onset response in serotonergic axons in A24b | - | - | 2697 | 11 | 4 |
| **1E** | Grating onset response in serotonergic axons in A24b | - | - | 1146 | 5 | 4 |
| **1F** | Visuomotor mismatch response in serotonergic axons in A24b | - | - | 1343 | 6 | 4 |
| **1G** | Fraction of serotonergic axons in A24b responsive to locomotion onsets | Hierarchical bootstrap | $<10^{-4}$ | - | **11** | 4 |
|  | Fraction of serotonergic axons in A24b responsive to grating onsets | Hierarchical bootstrap | 0.71 | - | **5** | 4 |
|  | Fraction of serotonergic axons in A24b responsive to mismatch onsets | Hierarchical bootstrap | 0.59 | - | **6** | 4 |
| | Fraction of serotonergic axons in A24b responsive to locomotion onsets vs grating onsets | Hierarchical bootstrap | $<10^{-4}$ | - | **11,5** | 4,4 |
|  | Fraction of serotonergic axons in A24b responsive to grating onsets vs mismatch onsets | Hierarchical bootstrap | 0.75 | - | **5,6** | 4,4 |
| | Fraction of serotonergic axons in A24b responsive to locomotion onsets vs mismatch onsets | Hierarchical bootstrap | $<10^{-4}$ | - | **11,6** | 4,4 |
| **1H** | Locomotion onset response in serotonergic axons in V1 | - | - | 1812 | 10 | 4 |
| **1I** | Grating onset response in serotonergic axons in V1 | - | - | 1050 | 5 | 4 |
| **1J** | Visuomotor mismatch response in serotonergic axons in V1 | - | - | 1159 | 6 | 4 |
| **1K** | Fraction of serotonergic axons in V1 responsive to locomotion onsets | Hierarchical bootstrap | $<10^{-4}$ | - | **10** | 4 |
| | Fraction of serotonergic axons in V1 responsive to grating onsets | Hierarchical bootstrap | $<10^{-4}$ | - | **5** | 4 |
|  | Fraction of serotonergic axons in V1 responsive to mismatch onsets | Hierarchical bootstrap | 0.88 | - | **6** | 4 |
| | Fraction of serotonergic axons in V1 responsive to locomotion onsets vs grating onsets | Hierarchical bootstrap | $<10^{-4}$ | - | **10,5** | 4,4 |
| | Fraction of serotonergic axons in V1 responsive to grating onsets vs mismatch onsets | Hierarchical bootstrap | $<10^{-4}$ | - | **5,6** | 4,4 |
| | Fraction of serotonergic axons in V1 responsive to locomotion onsets vs mismatch onsets | Hierarchical bootstrap | $<10^{-4}$ | - | **10,6** | 4,4 |
| **2B** | Correlation of serotonergic axon activity with locomotion velocity vs pupil diameter in A24b | Hierarchical bootstrap | 0.005 | **2697** | 11 | 4 |
| **2C** | Correlation of serotonergic axon activity with locomotion velocity vs pupil diameter in V1 | Hierarchical bootstrap | 0.03 | **1812** | 10 | 4 |
| **3A** | Mean activity of serotonergic axons in A24b in closed loop vs open loop | Hierarchical bootstrap | 0.18 | **351** | 3 | 3 |
| **3B** | Mean activity of serotonergic axons in V1 in closed loop vs open loop | Hierarchical bootstrap | 0.03 | **471** | 4 | 4 |
| **4A** | Mean activity of serotonergic axons in A24b in closed loop under low vs high visuomotor gain | Hierarchical bootstrap | 0.04 | **678** | 5 | 4 |
| **4B** | Mean activity of serotonergic axons in V1 in closed loop under low vs high visuomotor gain | Hierarchical bootstrap | $<10^{-4}$ | **232** | 3 | 4 |
| **S1A** | Locomotion onset response in serotonergic axons in A24b | - | - | 2697 | 11 | 4 |
| **S1B** | Locomotion onset response in serotonergic axons in V1 | - | - | 1812 | 10 | 4 |
| **S2** | Difference in locomotion onset response to grating onset response in serotonergic axons in A24b vs V1 | Hierarchical bootstrap | 0.03 | **1146,1050** | 5,5 | 4,4 |

